# Gene co-expression is distance-dependent in breast cancer

**DOI:** 10.1101/399253

**Authors:** Diana García-Cortés, Guillermo de Anda-Jáuregui, Cristóbal Fresno, Enrique Hernandez-Lemus, Jesús Espinal-Enríquez

**Affiliations:** Computational Genomics Division. National Institute of Genomic Medicine, México; Centro de Ciencias de la Complejidad, Universidad Nacional Autónoma de México

**Keywords:** Breast cancer molecular subtypes, loss of trans- regulation, Gene co-expression networks, Distance-dependent gene co-expression, Breast cancer networks

## Abstract

Breast carcinomas are characterized by anomalous gene regulatory programs. As is well known, gene expression programs are able to shape phenotypes. Hence, the understanding of gene co-expression may shed light on the underlying mechanisms behind the transcriptional regulatory programs affecting tumor development and evolution. For instance, in breast cancer, there is a clear loss of inter-chromosomal (trans-) co-expression, compared with healthy tissue. At the same time cis- (intra-chromosomal) interactions are favored in breast tumors. In order to have a deeper understanding of regulatory phenomena in cancer, here, we constructed Gene Co-expression Networks by using 848 RNA-seq whole-genome samples corresponding to the four breast cancer molecular subtypes, as well as healthy tissue. We quantify the cis-/trans- co-expression imbalance in all phenotypes. Additionally, we measured the association between co-expression and physical distance between genes, and characterized the proportion of intra/inter-cytoband interactions per phenotype. We confirmed loss of trans- co-expression in all molecular subtypes. We also observed that gene cisco-expression decays abruptly with distance in all tumors in contrast with healthy tissue. We observed co-expressed gene hotspots, that tend to be connected at cytoband regions, and coincide accurately with already known copy number altered regions, such as Chr17q12, or Chr8q24.3 for all subtypes. Our methodology recovered different alterations already reported for specific breast cancer subtypes, showing how co-expression network approaches might help to capture distinct events that modify the cell regulatory program.

## Background

A cellular context is determined by the coordinated expression of specific sets of genes, hence, gene transcription is a highly regulated process. In this sense, gene expression profiles and co-expression patterns might provide insight about shared transcriptional regulatory mechanisms [1, 2]. Among the elements involved in those regulatory mechanisms there are proteins that constitute the transcriptional machinery such as the RNA polymerase II and its associated enzymes [3], transcription factors and their cofactors, sequences identified by those transcription factors such as promoters [4] and enhancers [5, 6], histone modifications, both repressing or promoting gene expression, and structural proteins associated with chromatin architecture [7] (for a comprehensive review see [8, 9]).

Interactions among those elements allow the emergence of long-range regulatory instances and functional relationships have been characterized between regulatory elements and distal target genes [10]. Although these long-range instances are clearly important, transcription of a specific gene is also influenced by its physical location: gene order is not random and neighboring genes tend to be co-expressed in several eukaryotic cells [11–13]. Moreover, there is evidence that the transcription process contributes to the formation of gene clusters and influence chromatin organization at a fine scale [14].

In cancer there is a dramatic disruption of the transcriptional process leading to altered gene expression and the promotion of tumor progression [15]. Elements in nearly all levels of transcriptional control can suffer from modifications in cancer. A mutation can alter gene expression directly when it takes place in gene control sequences such as enhancers, promoters or insulators. On the other hand, transcriptional dysregulation can arise from genomic alterations indirectly when they modify the activity of signaling factors such as transcription factors, signaling proteins or when they cause a disruption in chromatin regulators or chromosome structuring proteins [16].

Heterogeneity in the aforementioned alterations is displayed in the diversity of cancer-associated gene expression profiles and their related phenotypes. In this sense, breast cancer and its molecular subtypes, Luminal A, Luminal B, HER2+ and Basal-like [17, 18], are an emblematic example because they have distinct patterns of gene co-expression profiles and lead to different breast cancer manifestations both at the molecular and clinical levels.

Breast cancer subtypes have been associated with different prognosis, survival rates and metastatic behavior [19, 20]. The longest survival has been observed in patients with the Luminal A subtype, followed by those with Luminal B and HER2+. Finally, Basal-like subtype patients have been associated with the poorest survival [21]. Median duration of survival from time of first distant metastasis follows the same trend [22]. Hence, outcomes in breast cancer can be traced back to the transcriptomic level.

The altered gene expression patterns observed in breast cancer are usually studied using next generation sequencing (NGS) techniques such as RNA-Seq [23]. Computational methods can be used to analyze the resulting data and to understand different features of the biological system. In particular, co-expression networks capture important aspects of the connectivity structure of the transcriptional profile [24, 25]. Moreover, global and local properties of co-expression networks may provide some insights into the actual biological mechanisms associated to the gene co-expression program [26–28].

Previously, by comparing the gene co-expression network derived from RNA-Seq data from the unclassified breast cancer samples in TCGA and the co-expression network from the healthy samples, we reported a significant reduction in the number of interactions linking inter-chromosomal genes (*trans*-) and an increase on the strenght of co-expresssion between intra-chromosomal (*cis*-) genes [29, 30]. On the other hand, in the healthy breast tissue, gene co-expression was apparently independent of the chromosome location of genes. This phenomenon evidenced that in breast cancer, there is an influence of the physical distance between genes and their related co-expression values.

With the aim to provide a broader and deeper landscape of the transcriptional regulation changes in the context of breast cancer heterogeneity, in this work, we characterized the *cis-/trans*- co-expression imbalance for each breast cancer molecular subtype compared with the co-expression profile of the healthy phenotype. We also assessed the gene co-expression distance dependency between *cis*- genes. Additionally, to have a more comprehensive description of the distance level of *cis*- co-expression, we characterized the *intra-cytoband/inter-cytoband* co-expression proportion per subtype.

With our network approach, we confirmed the *loss of trans- co-expression* in all breast cancer molecular subtypes. The *cis-/trans*- co-expression imbalance is specific for each subtype. We observed that gene *cis*- co-expression decays with distance in all breast cancer subtypes, but not in the healthy phenotype. We observed highly co-expressed gene hotspots, that tend to be confined at cytoband regions, and coincide accurately with already known copy number altered regions, such as the case for Chr17q12, or Chr8q24.3 for all subtypes.

## Results

### *cis-/trans*- imbalance is a common feature in breast cancer subtypes

To characterize the co-expression patterns for breast cancer molecular subtypes and the healthy phenotype, gene pair co-expression values were calculated (see methods section). Only the highest interactions in terms of their statistical significance (P ≤ 1*e*^−8^) were kept to better assess the differences among groups and the number of interactions for all datasets were reduced to match the lowest one (13,317 interactions, in the case of HER2+ co-expression network).

Fig. 1 exhibits circos plot representations of the most significant interactions for healthy and Basal subtype. Purple links represent *cis*- interactions, meanwhile orange links account for *trans*- relationships. Heatmap displays the number of adjacent links for every chromosome. Lowest row displays *cis*- interactions in purple, while the rest of the rows in the triangle refer to inter-chromosomal (orange) interactions. The number of *cis*- and *trans*- interactions for each phenotype is indicated in Table I and circos plot and heatmaps for all groups are included in Supplementary fig. 1.

**Figure 1:**
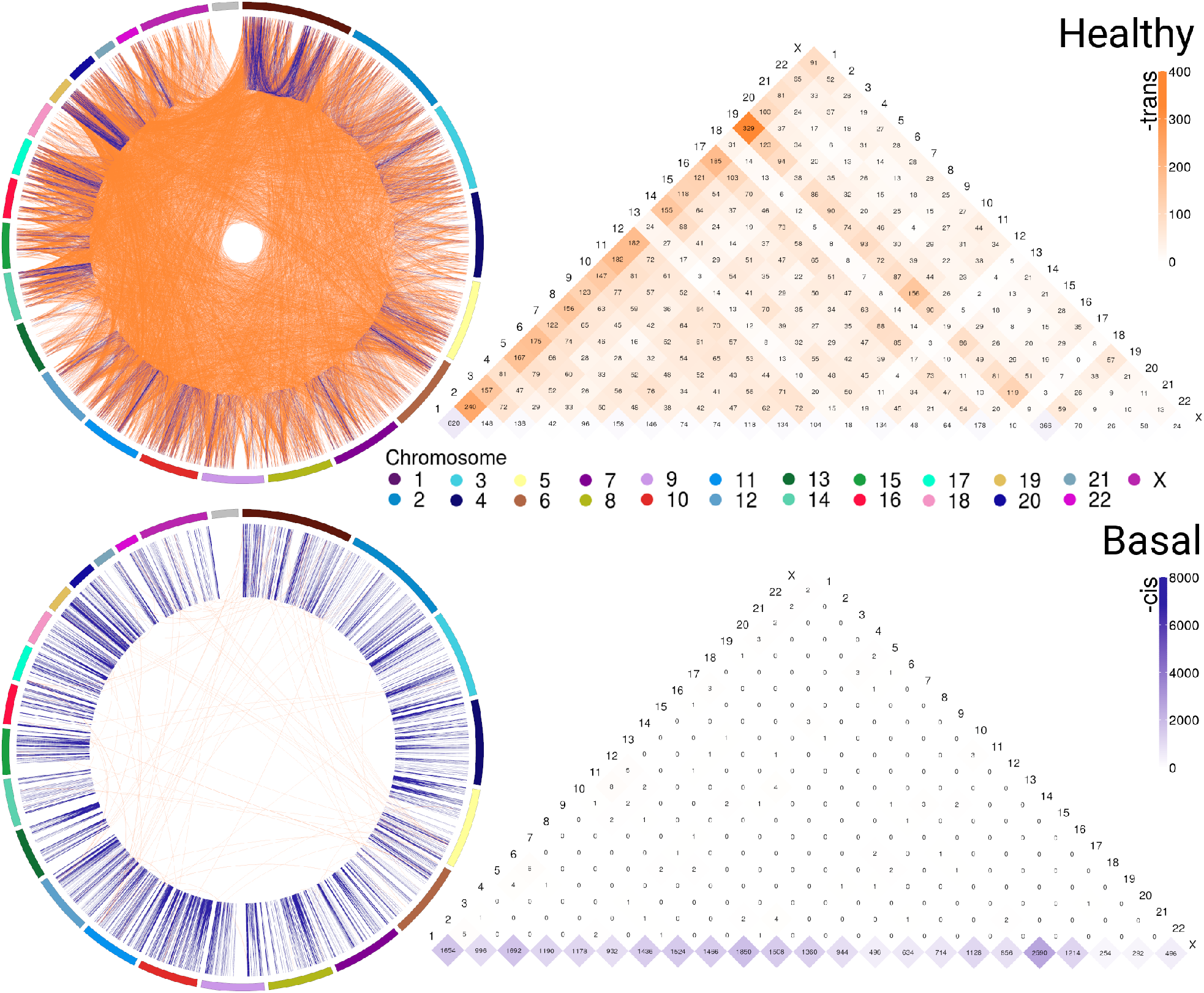
Abundance of *cis*-/*trans*- interactions in healthy and Basal networks. The circos plots and heatmaps show difference in *cis*-/*trans*- co-expression patterns between the healthy and Basal phenotypes. Plots contain the most significant interactions (P ≤ 1*e*^−8^). Orange links join genes from different chromosomes (*trans*-); meanwhile purple links represent *cis*- interactions. External arcs represent each chromosome. Heatmap shows the number of adjacent *cis*- and *trans*- links for both phenotypes in all chromosomes. Notice the difference in the purple/orange proportion. The Basal subtype is almost depleted of *trans*- (orange) interactions in all chromosomes.

**Table I:** Structural and distance-related features of the co-expression networks. Out of the 780 original breast cancer samples, 345 samples have been reliably classified into the molecular subtypes.

| Feature | Control | Breast cancer subtypes |  |  |  |
| --- | --- | --- | --- | --- | --- |
|  |  | Luminal A | Luminal B | HER2 | Basal |
| Number of samples | 113 | 217 | 192 | 221 | 105 |
| Number of genes | 4,922 | 4,843 | 5,156 | 5,092 | 5,262 |
| Total interactions | 13,317 | 13,317 | 13,317 | 13,317 | 13,317 |
| Intra (*cis*-) | 1,424 | 12,388 | 13,155 | 13,163 | 13,203 |
| Inter (*trans*-) | 11,893 | 929 | 162 | 154 | 114 |
| Clustering coeff. | 0.119 | 0.291 | 0.360 | 0.339 | 0.408 |
| *cis*-/*trans*- proportion | 0.120 | 13.335 | 81.204 | 85.474 | 115.816 |
| *cis*-/*trans*- proportion Z-score | 24.97 | 4,709.15 | 31,819.15 | 32,703.35 | 56,963.85 |
| Genes in LC | 4,005 | 1,200 | 174 | 154 | 120 |

The circos plot and the *cis-/trans*- proportion values in Table I shows that among the most significant co-expression relationships in the healhty phenotype there are more interactions between genes in different chromosomes than between genes in the same chromosome. Meanwhile, each one of the breast cancer molecular subtypes present an inverted proportion than the healthy phenotype indicating that *cis-/trans*- imbalance is a common feature in breast cancer subtypes. In fact, the Basal breast cancer heatmap shows that this subtype is almost depleted of *trans*- interactions. Additionally, *cis*- interactions in Basal circos plot appear to be straight lines, since genes with a high co-expression values are physically close.

The *cis-/trans*- proportion values where compared with those obtained from a configuration null model that takes into account the connectivity patterns displayed by the sets of the most significant co-expression interactions. According to Z-scores (Table I), *cis-/trans*- proportion values lie far from expected values in the configuration models, being the Basal suptype the set with the most extreme value. The fact that the healthy phenotype also displays a significant Z-score reveals that the *cis-/trans*- proportion in the healthy phenotype is greater even than what it is expected in a random model.

#### Gene co-expression networks show unique structures

Gene co-expression networks were assembled to analyze the structure that emerges from the most significant interactions. Fig. 2 displays the healthy network structure as well as the four network subtypes. In these visualizations, genes are colored according to the chromosome in which they are located. The five networks depicted show different structural features regarding size of components (set of nodes that are connected among them but have no interactions with other nodes) and gene connectivity.

**Figure 2:**
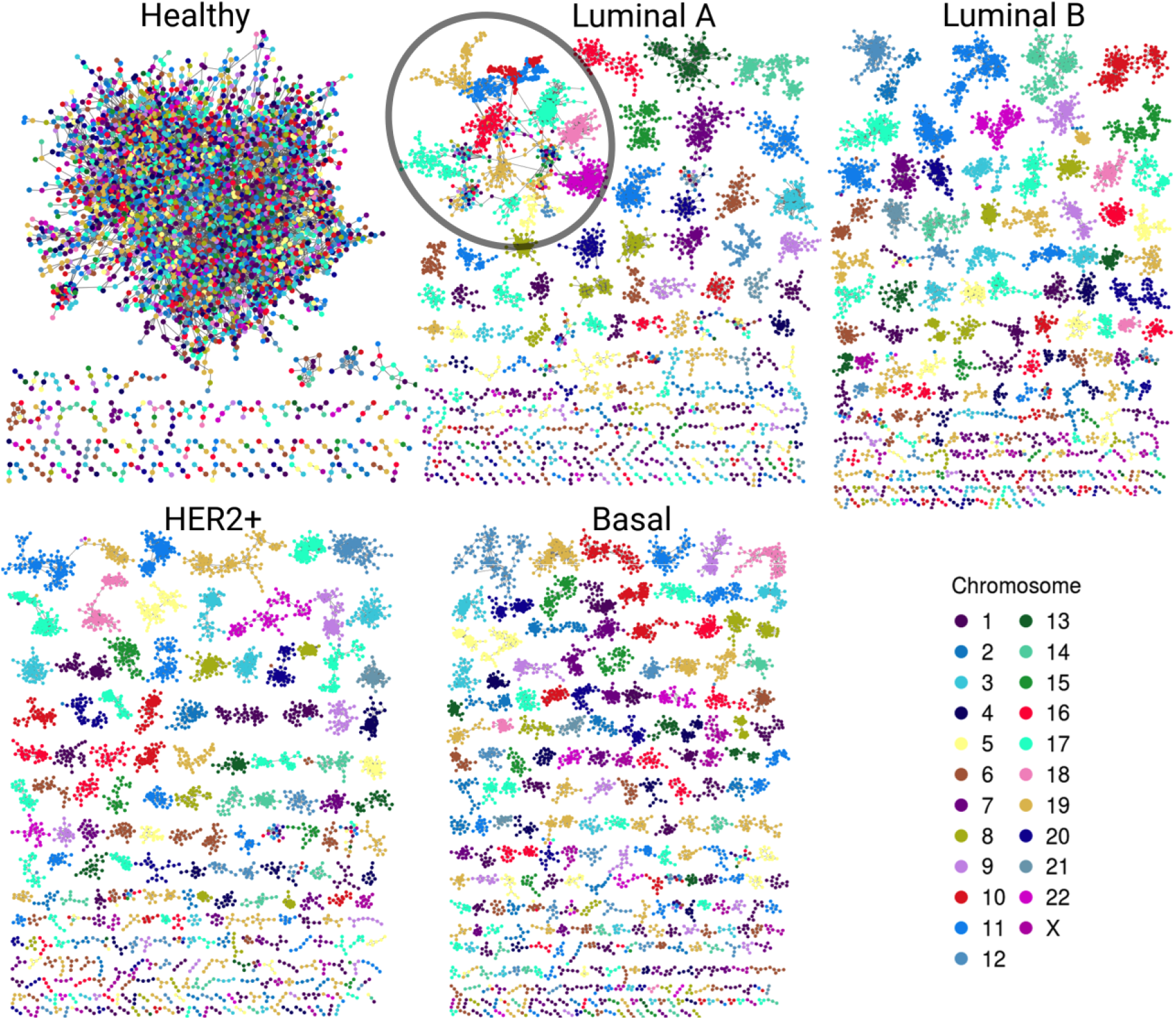
Gene co-expression networks. In this representation we observe the most significant interactions per network. Genes are colored according to the chromosome in which they are located. Notice the evident difference between size and color combination in the largest component in the healthy network versus the four subtypes. Cancer networks have almost the same color in the largest component. A particular case is the Luminal A subtype; despite largest component (circled) have several genes from different chromosomes, genes are grouped according to their corresponding chromosome.

In the healthy network, the largest component (LC) comprises almost all genes, contrary to any tumor subtype network, where the size of the LC is smaller (Table 1). Furthermore, the vast majority of genes in each component of any breast cancer subtype network belongs to a single chromosome. LC in the Luminal A subtype network displays a mixed behavior: while there are genes from different chromosomes linked inside the component, they are mostly assembled into clusters of *cis*- interactions. In terms of connectivity, genes in breast cancer networks are highly connected between them, i.e., network components in cancer networks present a higher clustering coefficient compared to the healthy network.

### Gene co-expression is distance-dependent in breast cancer

#### Physically close genes exhibit higher co-expression in breast cancer subtypes

To evaluate the relationship between gene-pair physical distance and the strength of co-expression, the entire set of *cis*- interactions were analyzed by calculating gene-pair co-expression values measured by Mutual Information (MI) for every *cis*- gene pair in the five phenotypes. MI values were normalized per phenotype to allow for comparison. In Fig. 3, gene pairs are grouped into 5 Mbp distance bins and the mean normalized-MI value for each bin is plotted. The area displays the standard deviation for each group.

**Figure 3:**
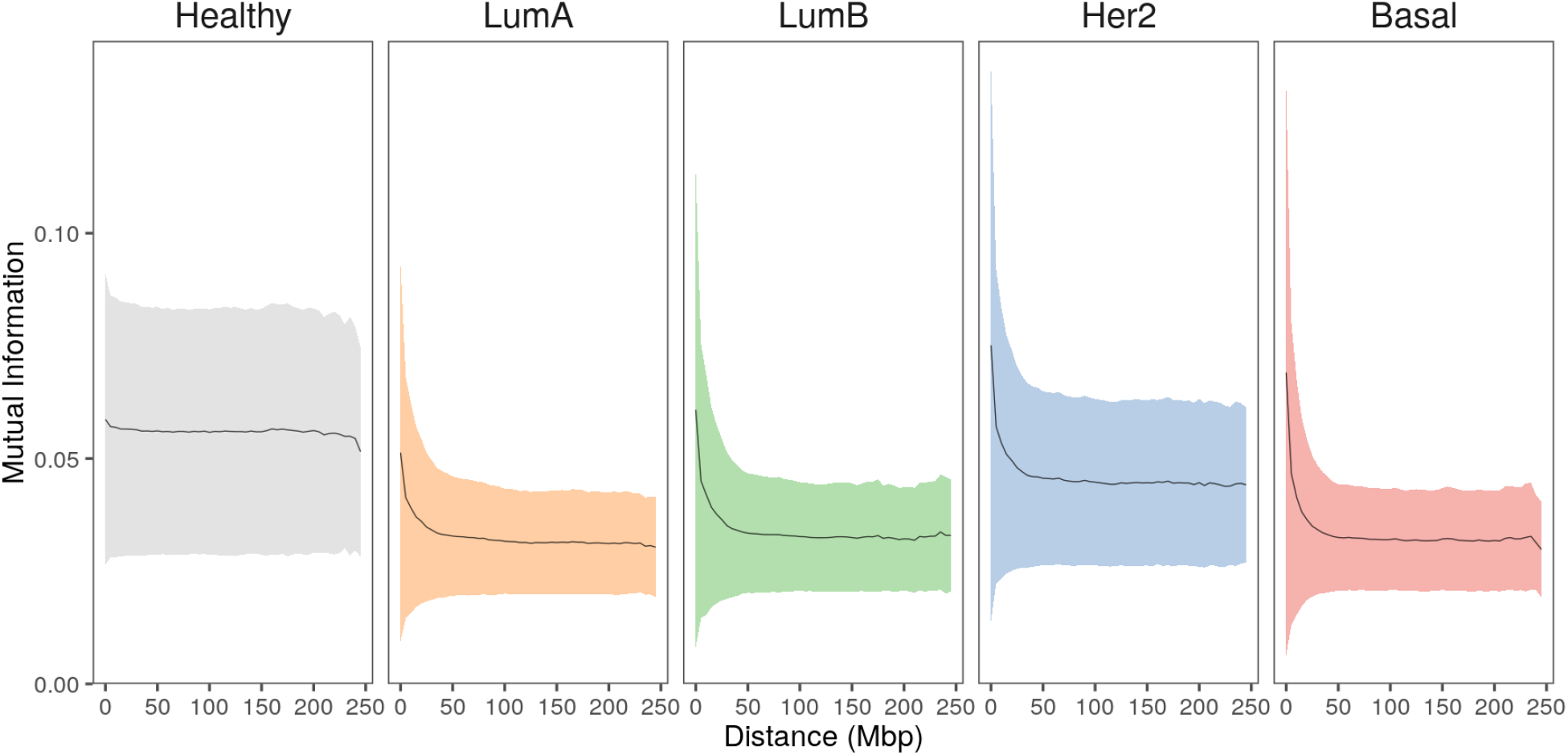
Strongest interactions occur between physically close genes in breast cancer subtypes. Mean co-expression MI values (black line) and the standard deviation (colored shadows) for all *cis*- interactions grouped at 5 Mbp gene-pair distance. In the case of healthy phenotype, co-expression remains constant at higher distances, contrary to any case of breast cancer. Furthermore, the plateau reached by all cancer phenotypes is clearly lower compared to the healthy one.

In the healthy phenotype plot, co-expression values remain almost constant with slightly greater MI values for the closest distances. Meanwhile, breast cancer subtype plots display similar behavior: short gene-pair distances have the highest MI values and, as distance increases, co-expression is lower, reaching a plateau close to 100 Mbp. Notice that the plateau in cancer is lower than the plateau observed in the healthy plot.

#### Co-expression decay associated to distance is specific for each chromosome and subtype

Once observed that gene co-expression decays with distance in each breast cancer subtype, the decay was evaluated for every chromosome individually. Normalized-MI values from *cis*- interactions were grouped into 100-gene-pair-bins and plotted versus gene-pair distance for every chromosome and for each subtype.

Fig. 4a displays plots for chromosomes 8 and 17 for the non-cancerous and the different breast cancer subtypes, while Fig. 4b displays values for all chromosomes in the healthy and the Basal phenotypes.

**Figure 4:**
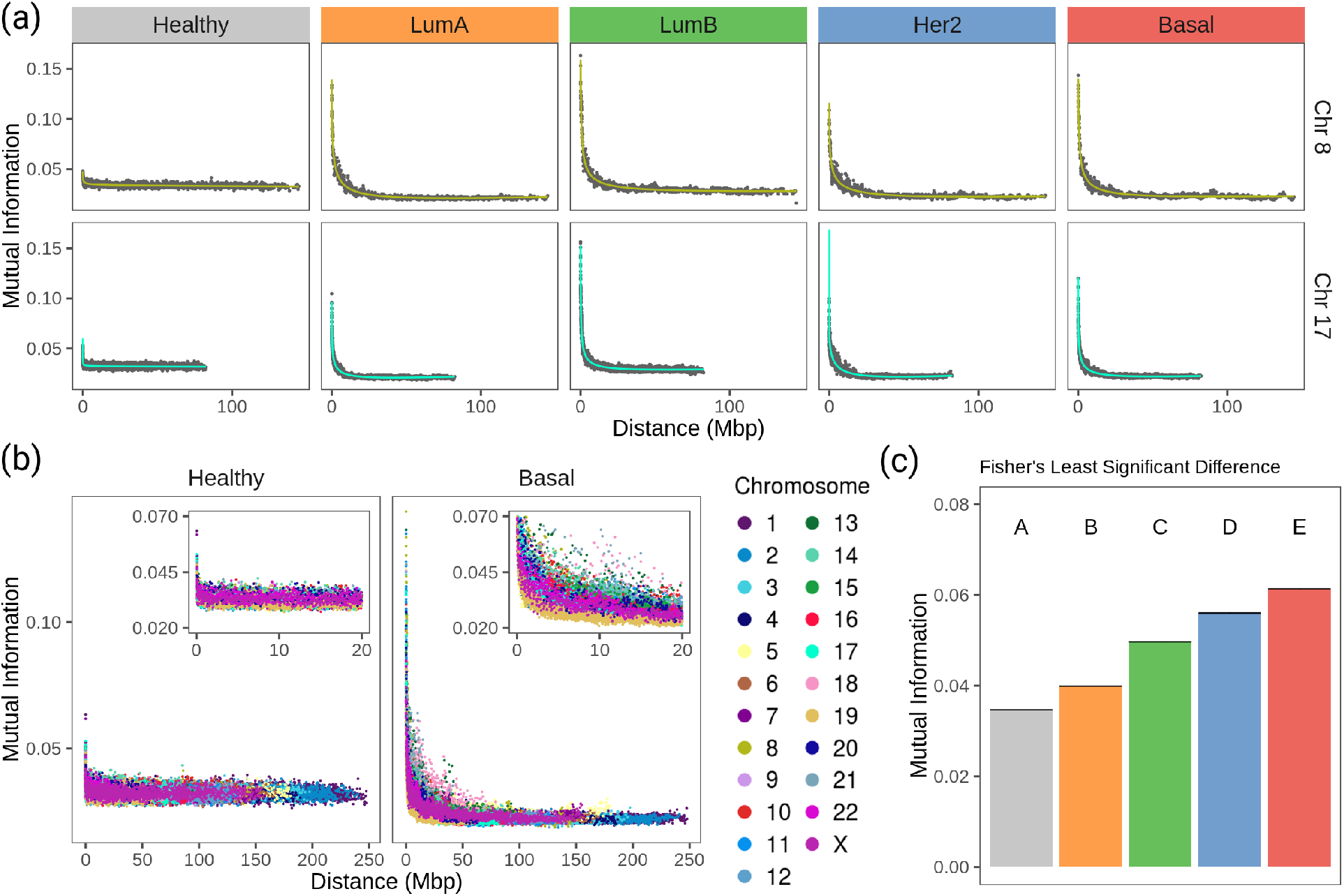
Strength of *cis*- insteractions decays with chromosomal distance in breast cancer subtypes. (a) Chromosomes 8 and 17 gene co-expression through distance. For both chromosomes, there is a remarkable difference between the strength of co-expression in healthy phenotype compared to any subtype. Each point represents the average of gene co-expression (MI) of 100 gene pair according to the distance between them. Green and turquoise curves are the fitting to a 4^*th*^ degree linear model. (b) Gene co-expression by distance in all chromosomes for healthy and Basal subtype. Color code of the chromosomes appears at the right of the figure. (c) Fisher’s Least Significant Difference test result of MI mean values at 1 Mbp gene-gene distance. Test assigns a different group for each phenotype, which confirms they are significantly different.

Values in the MI-distance plots were adjusted using a linear model that accounts for an independent intercept for each chromosome (see Methods section). The complete model has an adjusted R^2^ = 0.94. The ANOVA table results are presented in Supplementary Table 1, which confirms the distance-dependent decay in breast cancer subtypes (P <1e-4)). The shape of this decay in all cancer phenotypes is similar to that showed by our group for basal breast cancer [31]. There, co-expression strength diminished with distance in an exponential-like form. Supplementary fig. 2, contains the scatter plots for all chromosomes in the four breast cancer subtypes and the non-cancerous phenotype.

In Fig. 4a for any subtype, MI follows a decay relative to gene-pair distance. On the other hand, Fig. 4b shows that the strength of *cis*- interactions in the healthy phenotype is higher for extremely close gene pairs and then remains constant along the x-axis. Furthermore, the decay is different for each chromosome in breast cancer. This can also be observed in Fig. 4a, in Chr17 the decay is more abrupt than the case of Chr8.

Between chromosomes, the same subtype has slightly different patterns of decay, but that is not the case for the healthy co-expression program (Fig. 4). The inserts of Fig. 4b are a zoom over the closest distances. Color distribution allows the observation that there is more variability of *cis*- interactions in the Basal co-expression values than in the healthy phenotype.

By means of a Fisher’s Least Significant Difference test (see Methods section) it was proved that co-expression strength decay is different for each breast cancer subtype as they are classified into individual groups displayed in Fig. 4c.

### High density intra-cytoband hotspots emerge in breast cancer subtypes

Patterns of highly connected individual chromosome regions emerge for each subtype when individual chromosomes are analyzed. Cytogenetic bands where used to define chromosome regions of physically close genes: *cis*- interactions in the co-expression networks are classified into **intra-cytoband** when they link genes in the same cytogenetic band, and **inter-cytoband**, when genes are on different cytogenetic regions.

In the four subtype networks, a highly interconnected region composed only by *intra-cytoband* links is observed in chr8q24.3 (Fig. 5). This intra-cytoband hotspot is not recovered in the healthy network. This region is important because it is commonly amplified in breast cancer tumors associated with altered function of MYC [32]. Furthermore, intra-cytoband genes in p-arm of chromosome 8 are also highly connected in the four cancer networks.

**Figure 5:**
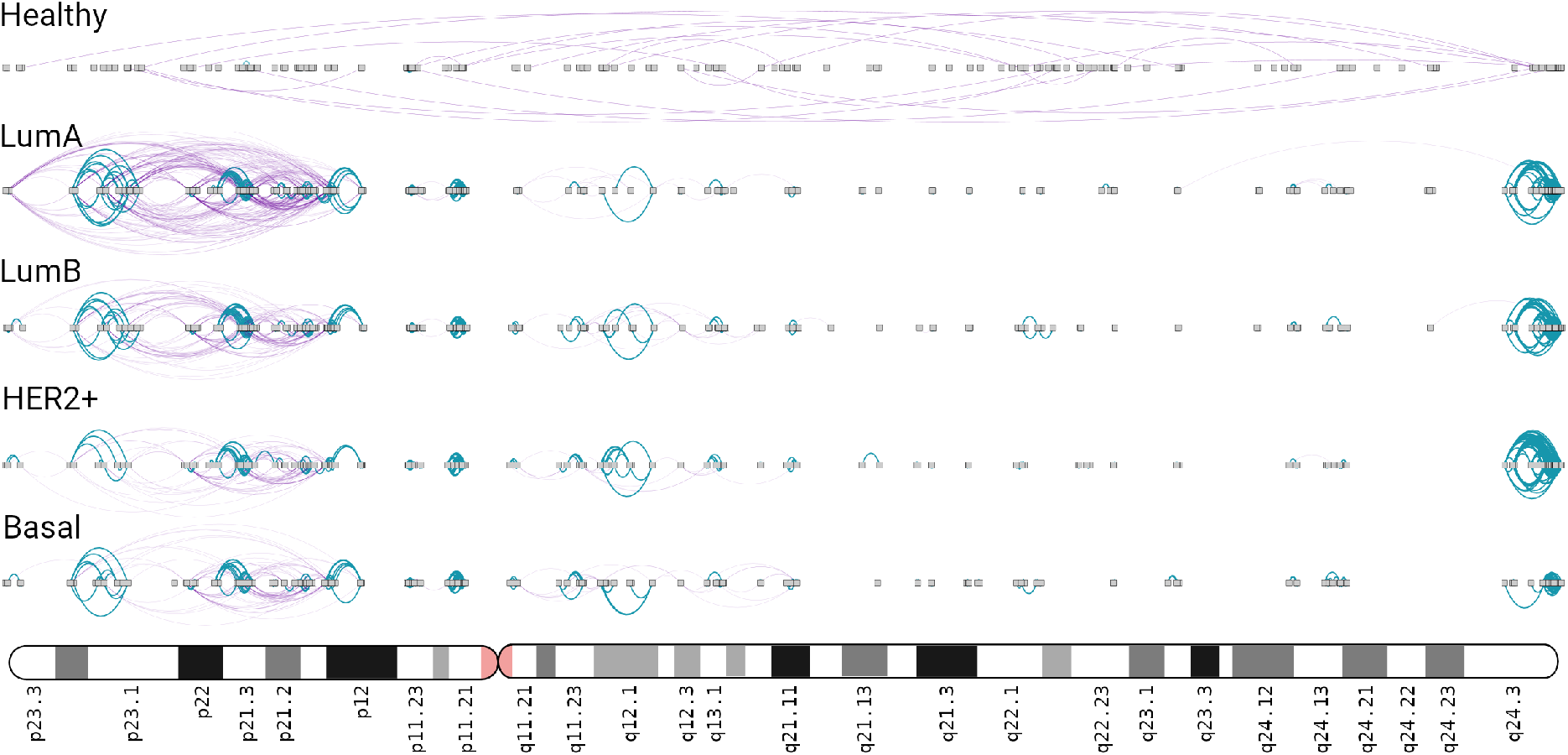
Chromosome 8 in breast cancer networks. This ideogram-like network layout shows the number of *intra-cytoband* (blue), *inter-cytoband* (purple), gene-gene interactions for all phenotypes. Notice the four hotspots observed in Chr8q24.3 region (right part) in the four subtypes, which is not the case of healthy network.

In Luminal A network, an important increment in *cis*- density in chromosome 17 is observed. A high number of *inter-cytoband* interactions in Chr17p can be appreciated (left part, Fig. 6). As in Figure 5, hotspots of co-expression can also be appreciated. p-arm of Luminal A network contains highly dense intra-cytoband interactions, but also inter-cytoband ones. In the rest of subtypes, intra-cytoband hotspots appear in different regions. Finally, chr17q25.3 region contains a highly dense hotspot in all subtypes. Bottom part of the figure is a zoom-in of chr17q12 region, where ERBB2 gene is located. There, several intra-cytoband interactions may be observed. The ideogram-like figures for all chromosomes in each phenotype are presented in Supplementary figs. 3 to 7.

**Figure 6:**
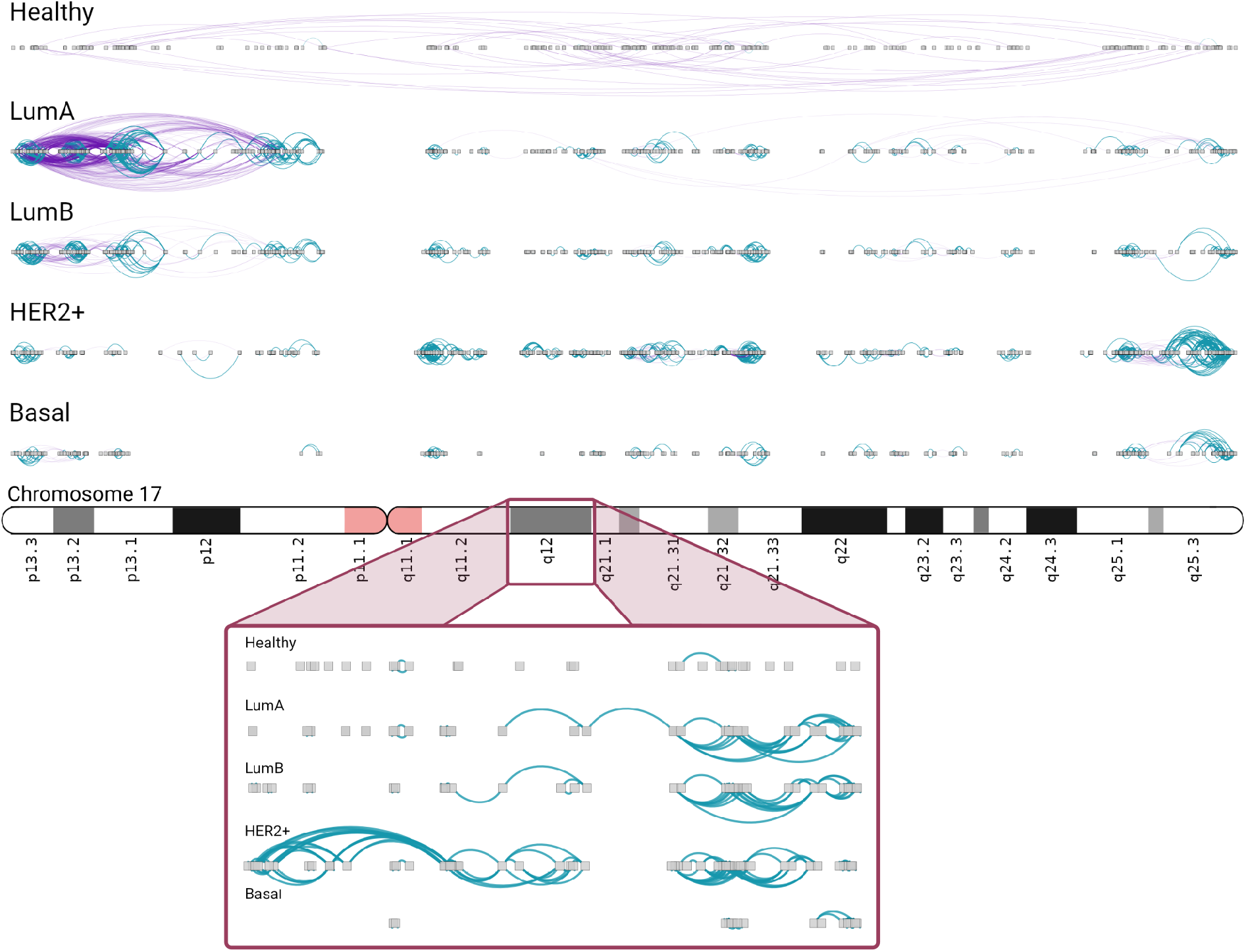
Chromosome 17 presents different *intra/inter cytoband* patterns for each subtype As in the previous figure, inter/intra-cytoband interactions are depicted for genes in chromosome 17. Notice the inversion of proportion from the healthy case to any other subtype. The bottom part corresponds to a zoom-in of chr17q12 region. Inter-cytoband links does not appear since we depicted one cytoband only.

### Strongest interactions are most commonly Intra-cytoband in breast cancer

To further analyze the co-expression patterns of the five phenotypes, we used cytogenetic bands since they delimit chromosome regions of physically close genes.

When genes linked by the most significant interactions are classified according to their cytogenetic band, the healthy phenotype displays a greater number of inter-cytoband than intra-cytoband links (apart from the already mentioned difference between *cis*- and *trans*- interactions). On the contrary, for any breast cancer subtype there are much more intra-cytoband interactions than inter-cytoband interactions.

In upper panel of figure 7 differences between intra-cytoband/inter-cytoband proportion between healthy and cancer networks are displayed. It is also remarkable the already mentioned difference between *cis*- and *trans*- interactions (orange bars). Boxplots in the bottom panel show the differences of co-expression values between intra/inter-cytobands, and also between *trans*- co-expression values.

**Figure 7:**
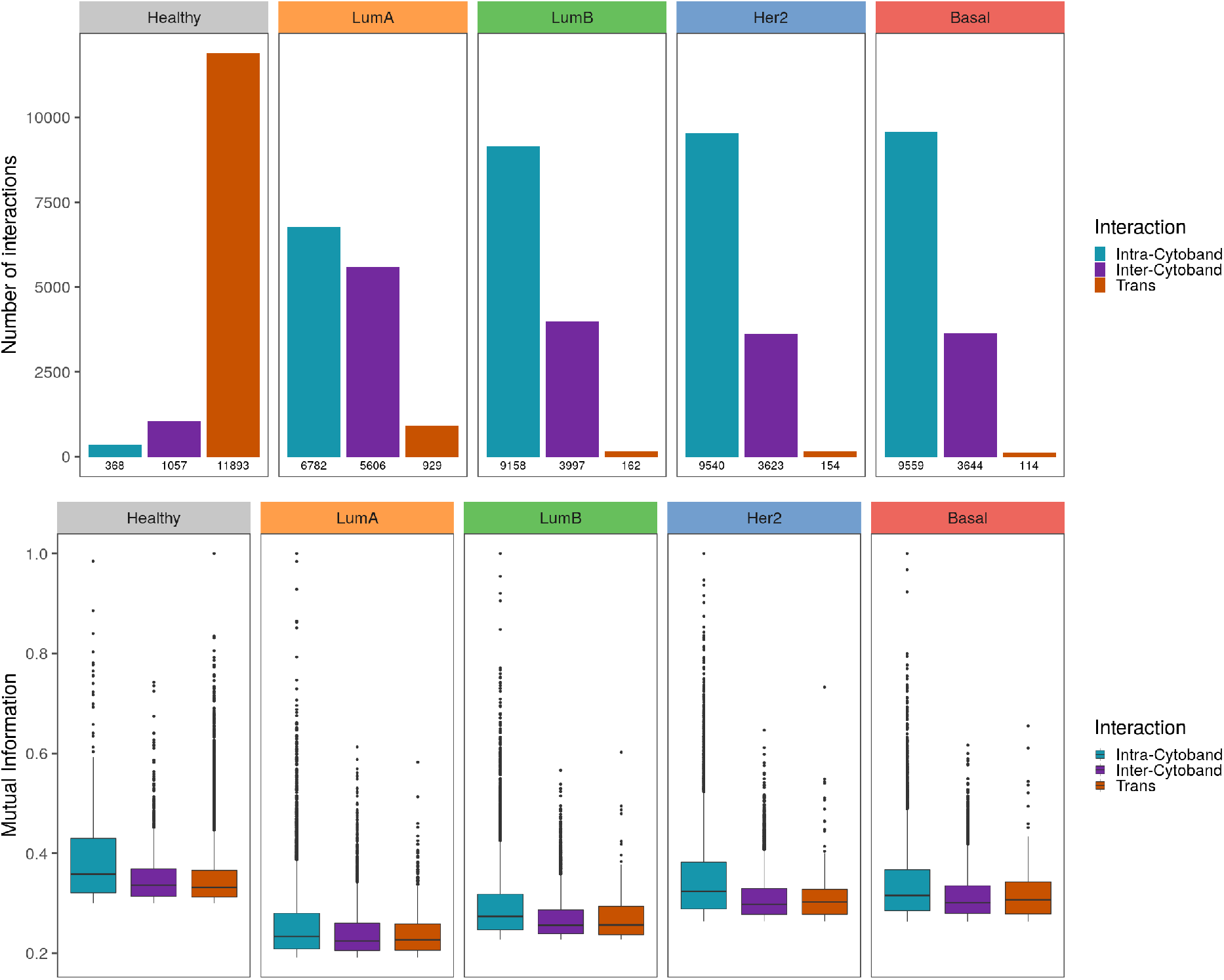
Strongest interactions are preferably Intra-cytoband in breast cancer Upper panel shows the number of *cis-intra-cytoband*, *inter-cytoband*, and *trans*- interactions for the five phenotypes. The lower panel shows boxplots of MI co-expression values for the same groups. To notice the higher MI values in healthy phenotype compared with the cancer ones.

## Discussion

In this work we have described the co-expression patterns in breast cancer molecular subtypes by analyzing the structure established by the strongest interactions in co-expression networks, as well as the relationship between the entire set of co-expression values and physical distance between gene pairs. There are four main characteristics displayed by the co-expression patterns in breast cancer molecular subtypes that are not present in the healthy phenotype: 1) there is a *cis*-/*trans*- interactions proportion imbalance, 2) the strength of gene-pair co-expression depends on physical distance, 3) there is an emergence of high density co-expression hotspots and 4) strongest interactions as well as hotspots are preferably intra-cytoband in breast cancer.

The *cis*-/*trans*- proportion imbalance in the co-expression networks is a phenomenon that was already observed for the unclassified set of breast cancer samples [29]. However, differences in transcriptional profile are widely accepted to recover clinical features such as prognosis and response to treatment [33]. Transcriptional profiles are the basis of the classification into the main four molecular subtypes: Luminal A, Luminal B, Her2-enriched and Basal. Our group has shown that co-expression networks of breast cancer molecular subtypes are different in terms of their modularity [34] and network architecture [35]. Here, it is shown that the *cis*-/*trans*- proportion imbalance is another feature that characterizes each network (Table I, Figure 3).

When compared to a null model, the healthy co-expression network received a significant Z-score on its *cis*-/*trans*- proportion. This is an expected observation given the well described influence from neighboring transcription in eukaryotic genome [12, 13]. Although it has been shown that some clustered genes are not only co-expressed but also functionally related [36], the phenomenon has been mostly explained as a mechanism to control dose-sensitive genes, allowing the modification of the expression of multiple genes by the same epigenetic marks [13, 37, 38].

In agreement with this feature, MI values for the healthy phenotype in Figure 3 display high co-expression at a very close distance but they quickly reach a plateau and then remain unchanged. In contrast, for any breast cancer subtype, there is a decay in gene co-expression associated with physical gene pair distance. This result suggests that co-expression from neighboring genes gets exacerbated in breast cancer and presents different patters for each molecular subtype.

During the oncogenic process, there is a disruption of the regulatory elements that control gene transcription [15]. The result is an altered expression profile associated with the promotion of a tumorigenic phenotype. Among these alterations, there are some examples that might favor the formation of clustered genes with similar expression. For instance, it has been shown that Topologically Associated Domains (TAD) boundaries are altered in cancer promoting the formation of additional TADs, smaller and within normal TAD-architecture [39, 40].

Furthermore, Estrogen receptor, usually upregulated in the luminal subtypes, is known to have an influence in the structural organization of chromatin in estrogen-regulated genes, particularly, by promoting long range interactions [41–44]. This feature may have an influence in the co-expression profiles, for example, in the higher number of *trans*- interactions in Luminal subtypes (Figure 2). Additionally, Luminal A and Luminal B maximal co-expression values at closest distances are smaller than in the remaining breast cancer subtypes (Figure 3).

Alterations in the methylation patterns of contiguous gene regions have also been identified in cancer. Through hypermethylation and subsequent gene repression, regions of long-range epigenetic silencing (LRES) are found to be functionally inactivated in breast [45] and prostate cancer [46]. Long-range epigenetic activation (LREA) of adjacent genes within a region have also been found in prostate cancer [47]. These alterations might favor the formation of high density intra-cytoband hotspots, observed in the breast cancer co-expression profiles but not in the healthy one.

Among the most documented genomic modifications in breast cancer are gene copy number alterations (CNAs). CNAs are also known to affect gene expression over clusters of genes [48–50]. Amplification and protein overexpression of HER2 actually define a specific molecular subtype [51, 52]. According to [53], the HER2-amplicon was narrowed to an 85.92 kbp region including the TCAP, PNMT, PERLD1, HER2, C17orf37 and GRB7 genes. Four out of those six genes appear in the HER+ co-expression network, HER2 included.

There is another amplicon that has been associated with the BRCA1 mutated triple negative breast cancer in the region 17q25.3, that was also observed in HER2+ and Luminal B subtype [54] (Figure 6). We recovered the region as a intra-cytoband hotspot but it was more dense in the HER2+ subtype. Region 6p21.2-6p12 in the triple-negative breast cancer, associated with the Basal subtype has been also identified as an amplicon [55] and as a hotspot in our network (Supplementary fig. 7). We also identified the 17q21.3 amplicon in the HER2+ subtype [49].

Chr 16p13 hotspot was also found in our networks for all cancer phenotypes, but not for the healthy one. Genomic loss in this region has been associated with poor prognosis in colorectal cancer [56]. However, it has been observed a copy number gain in breast cancer [57]. Here we report a co-expression hotspot in said region, which coincides with the observation of [57]. Hence, the observed hotspots may serve as a proxy of other genomic and epigenomic alterations in cancer.

Alterations are not isolated. In addition to the already mentioned genomic modifications observed in cancer, it is known that the three-dimensional chromatin organization influences chromosomal alterations at the sequence level [58]. Hence, there is an interplay between the different regulatory elements altered during the oncogenic process. However, all of these changes are reflected in the gene expression profile.

Despite the lack of an accurate experimental setup to assess these results, our methodology recovered different instances of alterations that were already reported for specific breast cancer subtypes. This is an example of how a co-expression network approach might help to understand the different layers of transcriptional regulation that promote a tumorigenic phenotype.

## Conclusions

The heterogenic nature of breast cancer makes it a highly complex disease. The only way towards a profound comprehension of its functioning must be addressed from all possible perspectives. In this paper, we have used a co-expression network approach to dissect with high accuracy, the dependency of physical distance on gene co-expression in cancer. Several chromosome regions displaying a subtype-specific co-expression pattern coincide with genomic and epigenomic alterations already observed in breast cancer using different experimental protocols, as well as distinct -omic approaches.

From this work we highlight the following: *i)* gene co-expression networks capture different instances of molecular alterations in breast cancer, which can be translated to any other tissue. *ii)* This is a different approach to analyze alterations at the transcriptional level in cancer, and *iii)* The loss of long-range co-expression, concomitant with the strong gain of physically close gene co-expression, may constitute a manifestation of a non-previously observed phenomenon in cancer, and it can be the starting point to conduct investigations on how the transcriptional process shapes the oncogenic phenotype.

## Methods

### Databases

A collection of The Cancer Genome Atlas (TCGA) breast invasive carcinoma datasets were used in this work [26]. Briefly, 780 tumour and 101 normal IlluminaHiSeq RNASeq samples were adquired and pre-processed to *log*_2_ normalized gene expression values were obtained as described in [29].

### Data processing

The tumor *log*_2_ normalized expression values were classified using PAM50 algorithm into the respective intrinsic breast cancer subtypes (Normal-like, Luminal A, Luminal B, Basal and HER2-Enriched) using the Permutation-Based Confidence for Molecular Classification [59] as implemented in the *pbcmc* R package [60]. Tumour samples with a non-reliable breast cancer subtype call, were removed from the analysis as described in Table II.

**Table II:** Breast samples used in this work Notice that out of the 780 original breast cancer samples only 345 samples have been reliably classified into the molecular subtypes.

|  | **Control** | **Luminal** |  | **HER2+** | **Basal** |
| --- | --- | --- | --- | --- | --- |
|  |  | **B** | **A** |  |  |
| **Number of samples** | 101 | 58 | 163 | 72 | 142 |

Multidimensional Principal Component Analysis (PCA) over gene expression values showed a blurred overlapped pattern among the different breast cancer subtypes. Hence, multidimensional noise reduction using ARSyN R implementation was used [61]. Finally, PCA visual exploration showed that the noisy pattern was removed, thus breast cancer subtypes clustered without overlap.

#### Statistical analyses

Fisher’s least significant difference analysis (FLSDA) was performed to test whether MI distributions in the different groups were significantly different or there is some overlapping. FLSDA is a two-stage test for (multiple) pairwise comparison. The first stage consists in performing a “global” test for the null hypothesis that the expected value of the mean of the different groups is equal. In the (trivial) case that this null hypothesis is accepted, no further analysis is needed. If the global null hypothesis is rejected at a pre-specified level of significance, then a second stage is performed on all pairwise comparisons at the same level of significance [62]. Here we performed FLSDA with the LSD.test function in R using the default significance value (*α* = 0.05) and FDR correcting for multiple testing.

#### Network Construction

Gene regulatory network deconvolution from experimental data has been extensively used to unveil co-regulatory interactions between genes by looking out for patterns in their experimentally-measured mRNA expression levels. A number of correlation measures have been used to deconvolute transcriptional interaction networks based on the inference of the corresponding statistical dependency structure in the associated gene expression patterns [63–67]. It has long been known that the maximum likelihood estimator of statistical dependency is mutual information (MI) [66–69]. ARACNE [70] is the flagship algorithm used to quantify the degree of statistical dependence between pairs of genes.

In a nutshell, the algorithm calculates the *Mutual Information (MI)* –a non-parametric measure that captures non-linear dependencies between variables– in a relatively fast implementation. The method associates a *MI* value to each significance value (p-value) based on permutation analysis, as a function of the sample size. To make comparable the five networks, the highest interactions in terms of their statistical significance (P ≤ 1*e*^−8^) were kept. In this way, we better assessed the differences among groups. The number of interactions in the five networks were reduced to 13,317 interactions, which is the lowest number of interactions in our networks with significant p-value. This was the case of HER2+ network.

## Abbreviations

LC: Largest Component
MI: Mutual Information
TCGA: The Cancer Genome Atlas

## Availability of data and materials

The datasets analysed during the current study are available in the GDC Legacy Archive repository, https://portal.gdc.cancer.gov/legacy-archive.

## Author’s contributions

DGC performed computational analyses, developed and implemented programming code, performed pre-processing and low-level data analysis, made the figures, drafted the manuscript. GDJ developed and implemented programming code, contributed to the writing of the manuscript. CF performed pre-processing and low-level data analysis, contributed to the writing of the manuscript. EHL devised the project’s strategy and methodological approach, contributed to the theoretical and modeling analysis, co-supervised the project, contributed and supervised the writing of the manuscript. JEE conceived and designed the overall project, co-supervised the project, took the lead in the biological analyses, drafted the manuscript. All authors read and approved the final version of the manuscript.

## Supplementary Material

**Supplementary Fig. 1**. Circos plots contain the 0.01% highest mutual information values for each network. Orange lines join genes from different chromosomes; meanwhile the blue lines take account for intra-chromosomal (*cis*-) interactions. External circles indicate the chromosome band. Bottom right panel is a heatmap representing the number of *cis*- and *trans*- interactions per chromosome for each subtype.

**Supplementary Fig. 2**. Scatter plots showing the strength of *cis*- interactions with respect to physical distance in all chromosomes for the five phenotypes.

**Supplementary Fig. 3**. *cis*- interactions for healthy network. Blue edges are *intra cytoband*, meanwhile *inter cytoband* interactions are colored in purple.

**Supplementary Fig. 4**. *cis*- interactions for Luminal A network.

**Supplementary Fig. 5**. *cis*- interactions for Luminal B network.

**Supplementary Fig. 6**. *cis*- interactions for HER2+ network.

**Supplementary Fig. 7**. *cis*- interactions for Basal network.

**Supplementary Table 1**. Type-III ANOVA table results for the Intra-chromosomal regulation decay model. The ANOVA table tested the dependent of the Mutual Information (MI) values using a linear model including a fourth-order polynomial for the distance (D) between gene-pairs, accounting independent intercepts for each chromosome (Chr), breast cancer subtype and also for their corresponding polynomial terms. The model was fitted in the natural logarithm (Ln) scale for both MI and D. Significance codes: 0 ‘***’ 0.001 ‘**’ 0.01 ‘*’ 0.05 ‘.’ 0.1 ‘ ’ 1.

