## Supplementary materials file for "Gene co-expression is distance-dependent in breast cancer"

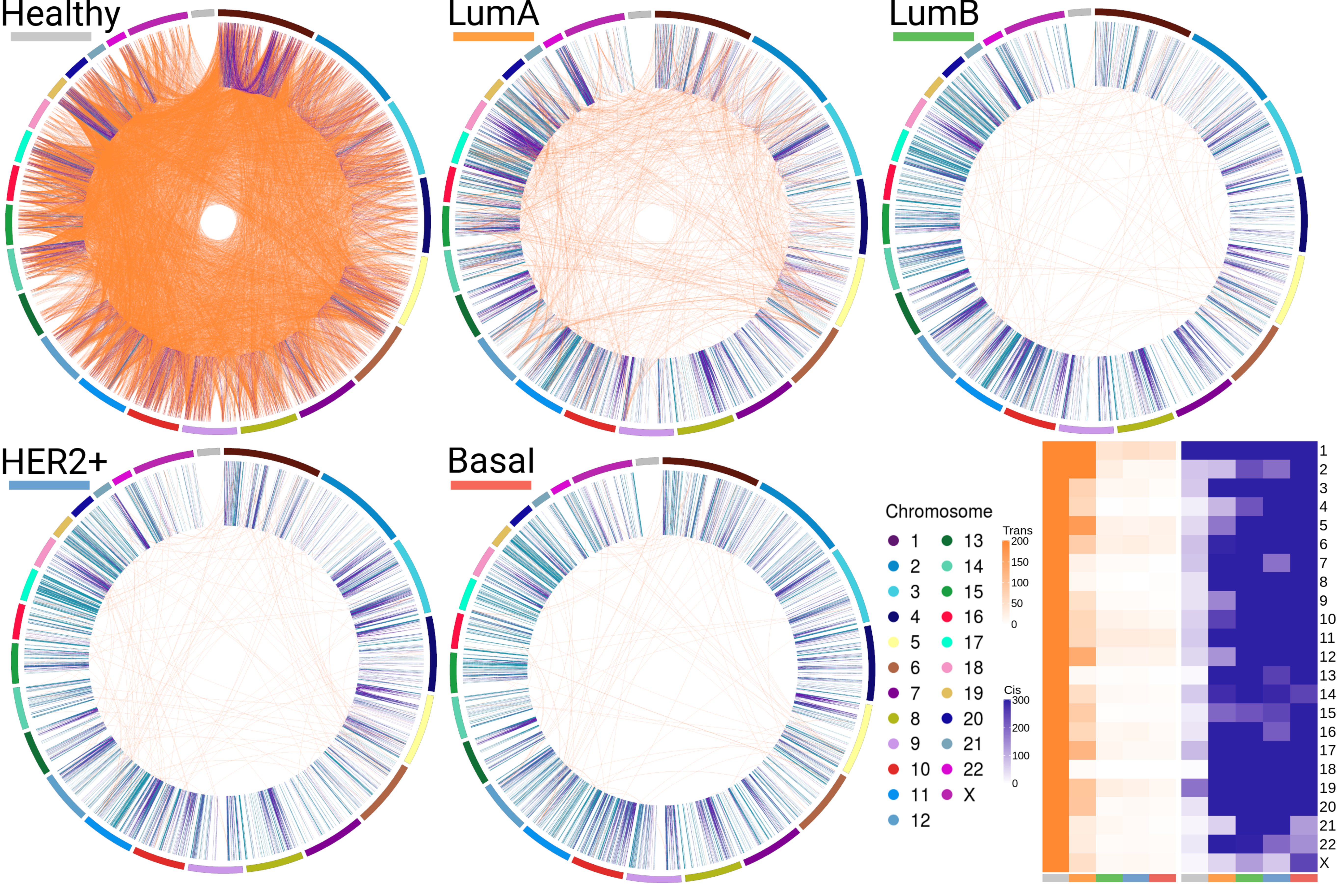

Healthy
LumA
LumB
HER2+
Basal
Chromosome
X
Trans
200
150
100
50
0
Cis
300
200
100
0

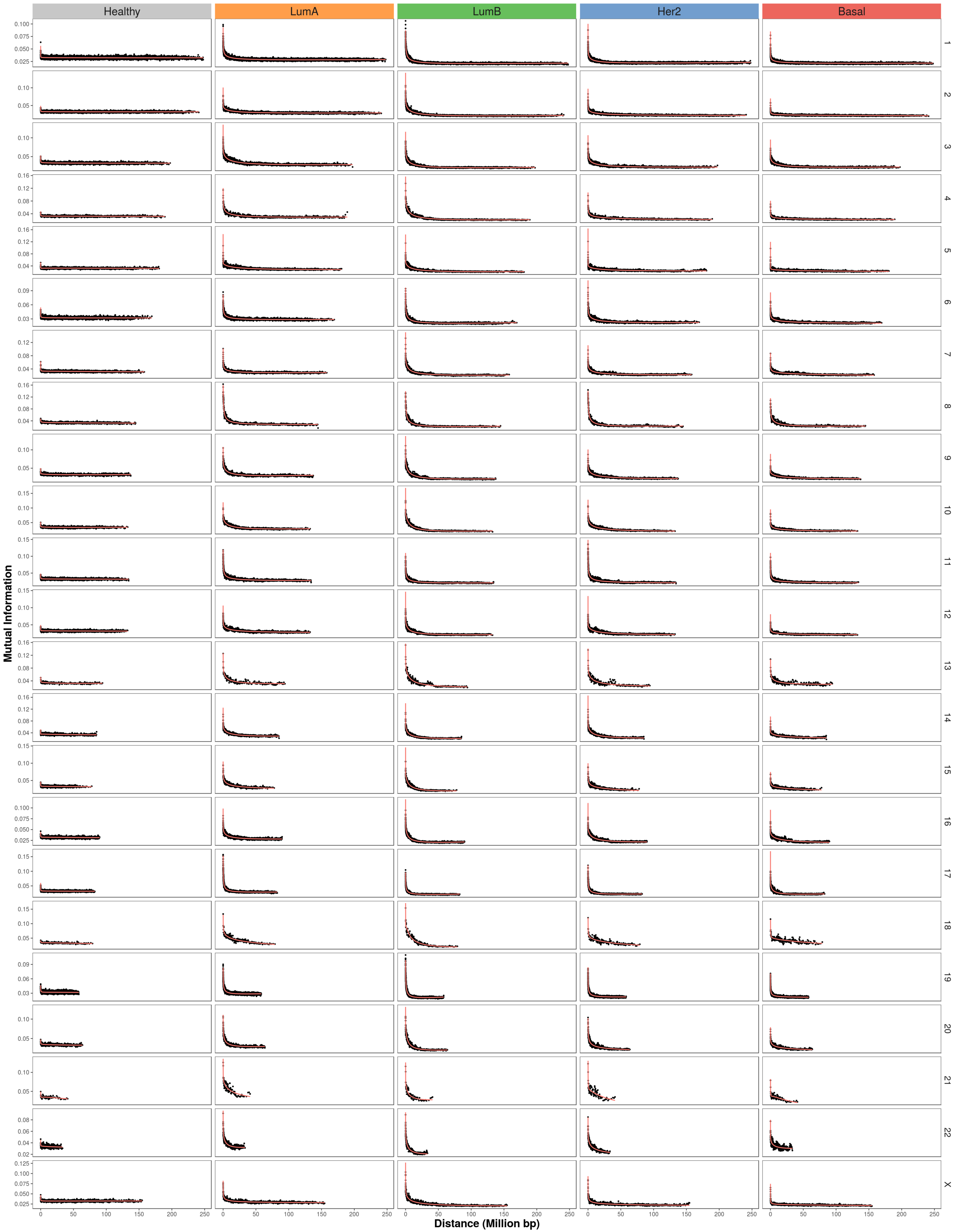

Healthy
LumA
LumB
Her2
Basal
X
Mutual Information
Distance (Million bp)
0
50
100
150
200
250

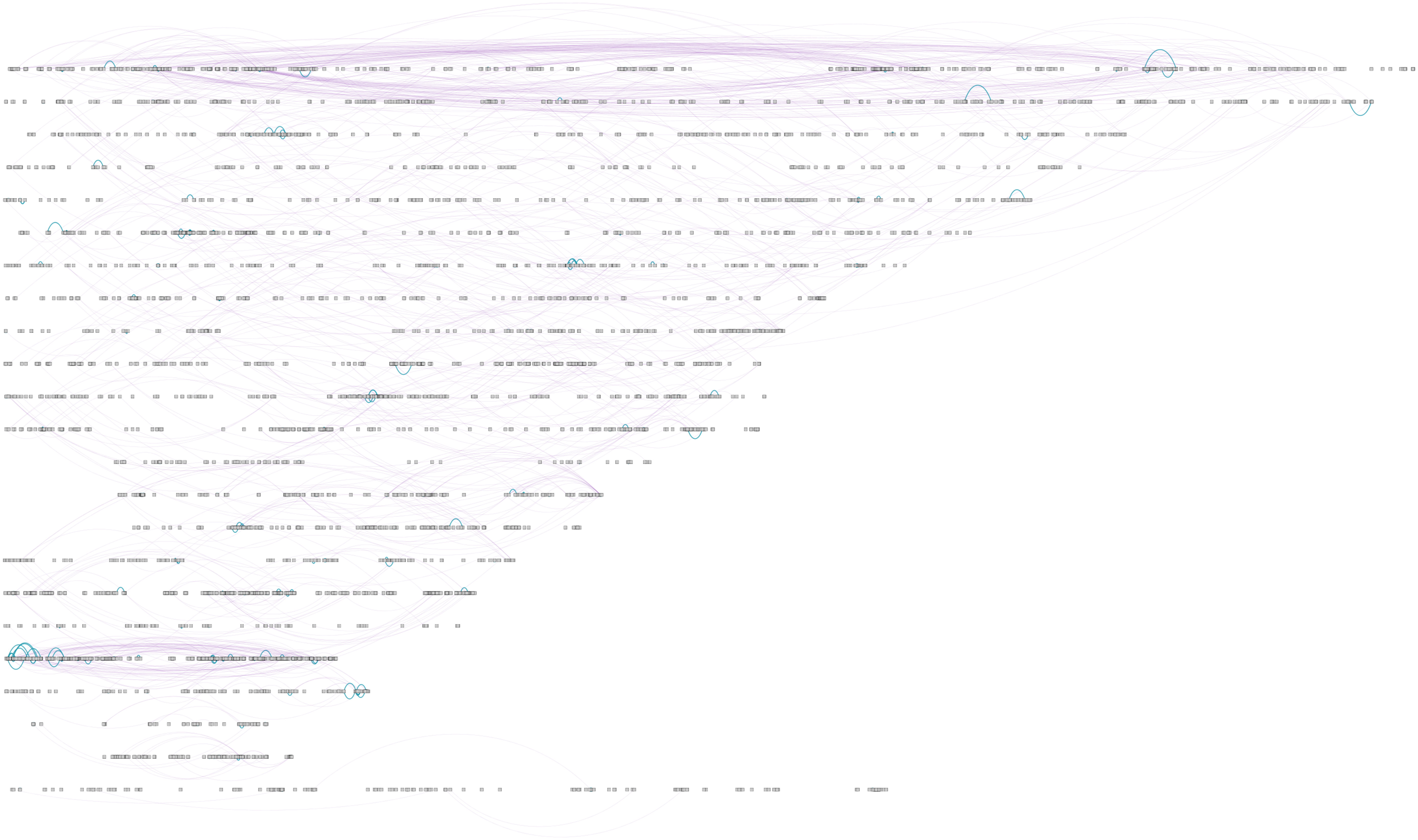

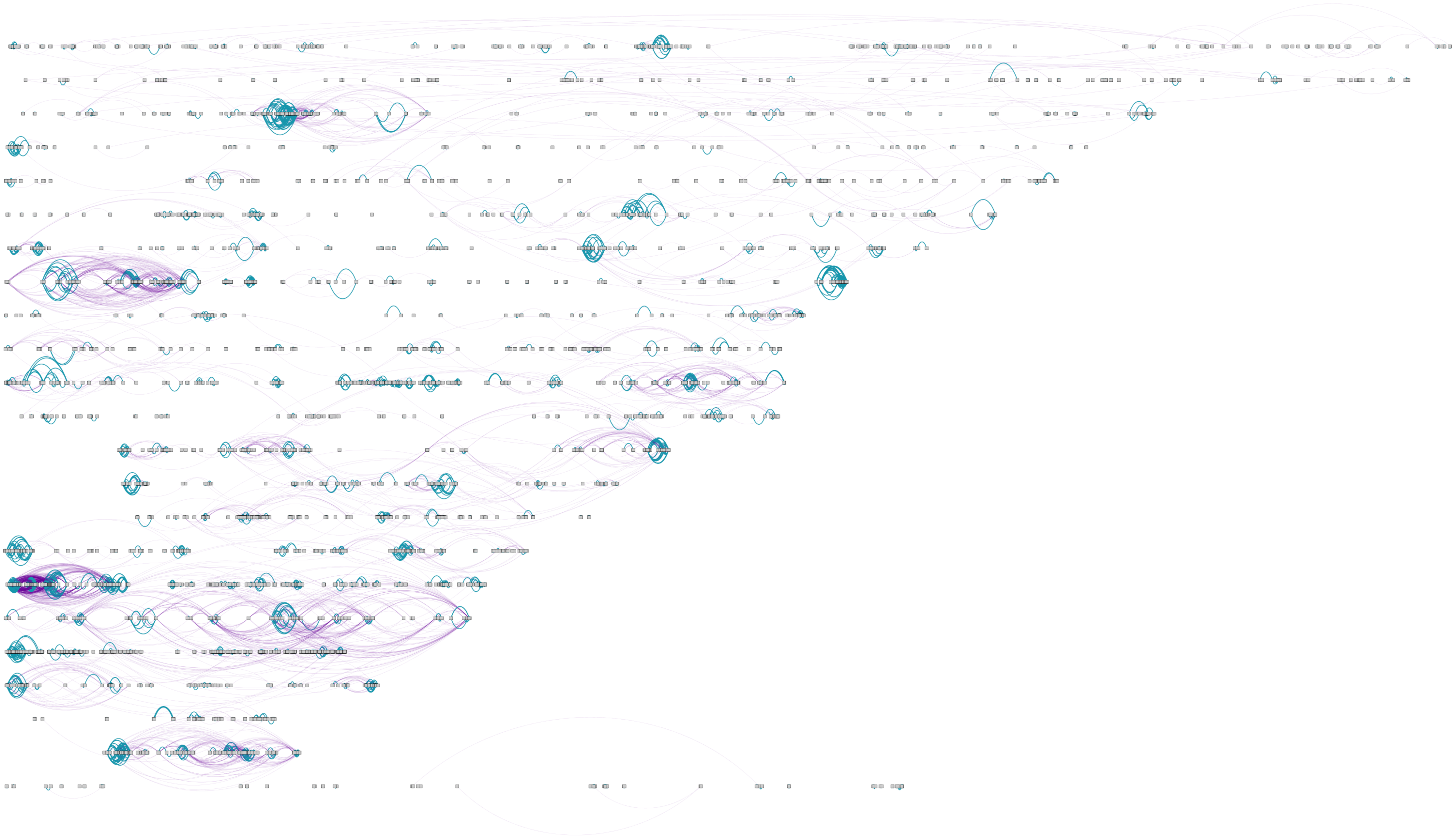

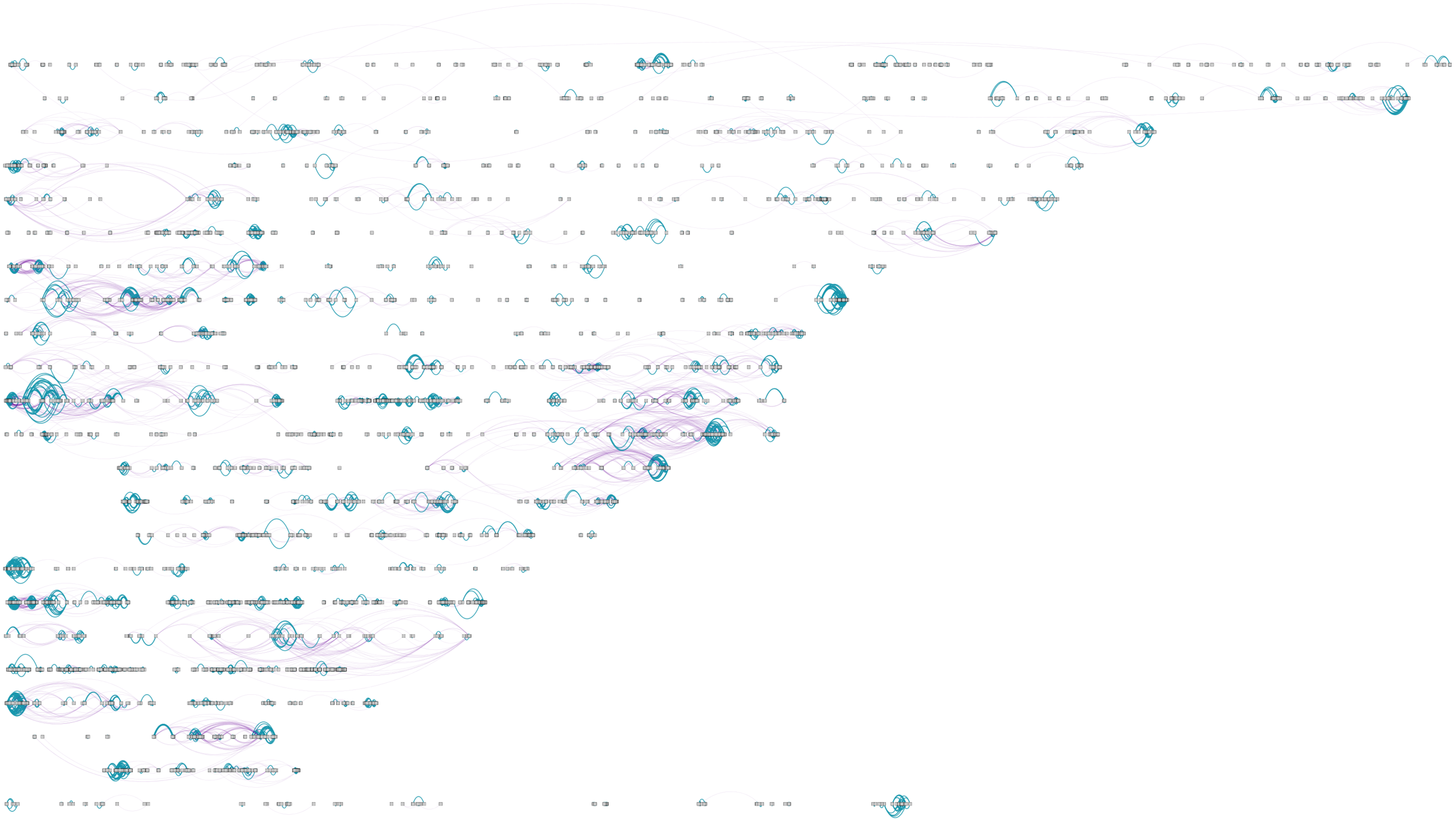

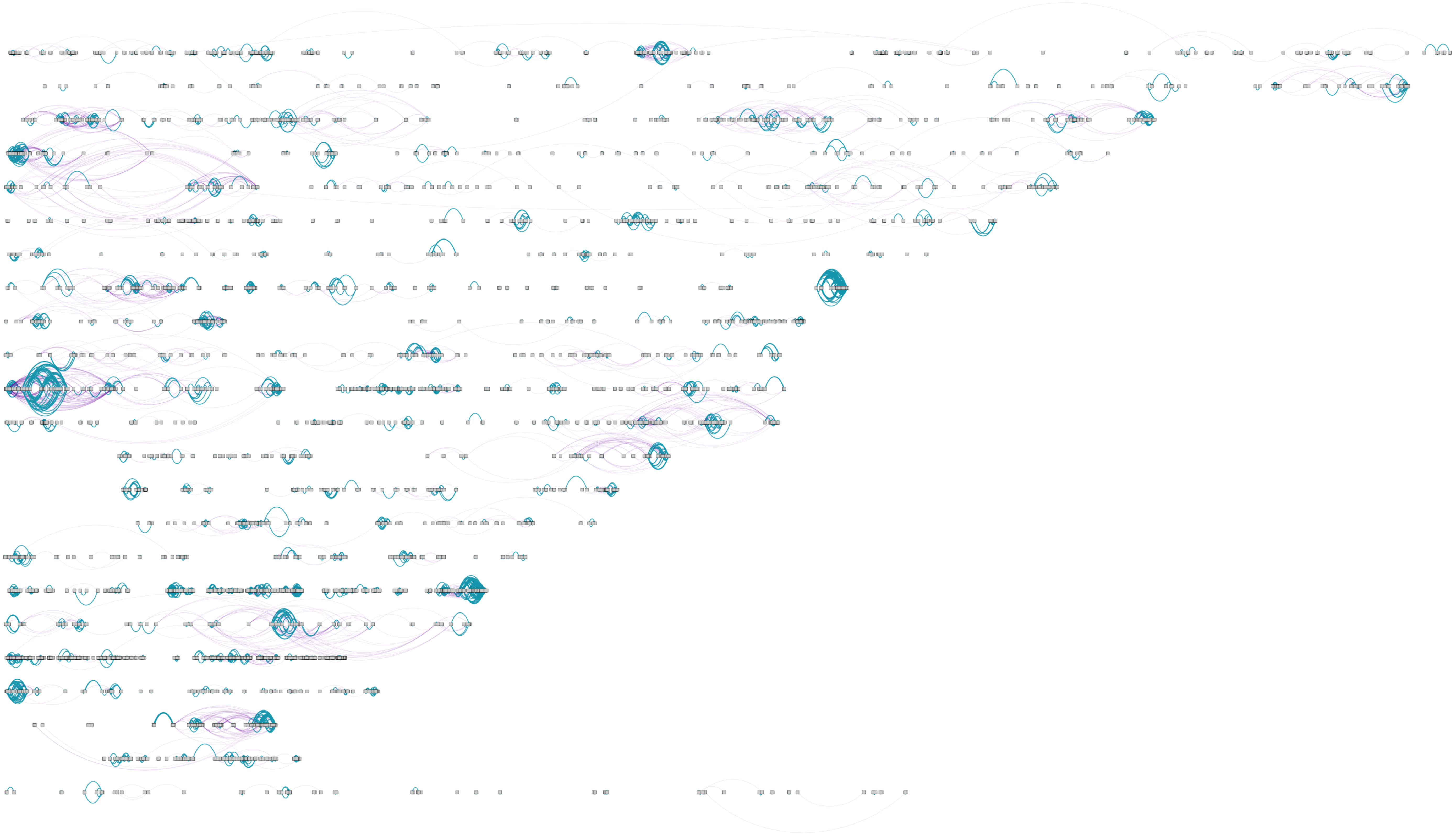

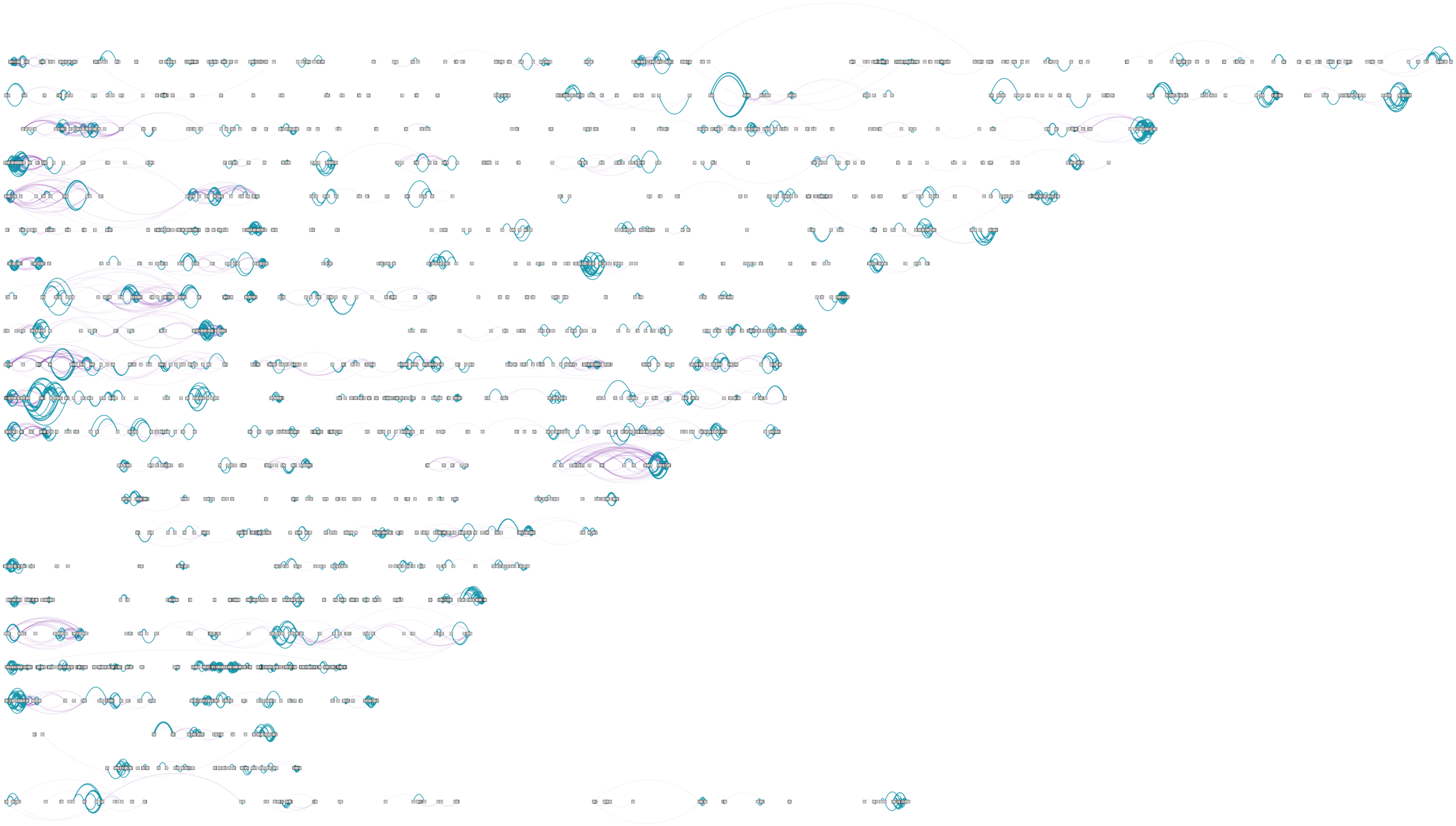

### Table S1

Table 1: Type-III ANOVA table results for the Intra-chromosomal regulation decay model. The ANOVA table tested the dependent of the Mutual Information (MI) values using a linear model including a fourth-order polynomial for the distance (D) between gene-pairs, accounting independent intercepts for each chromosome (Chr), breast cancer subtype and also for their corresponding polynomial terms. The model was fitted in the natural logarithm (Ln) scale for both MI and D. Significance codes: 0 '***' 0.001 '**' 0.01 '*' 0.05 '.' 0.1 ' ' 1.

| Model effect | Sum of Squares | Degree of Freedom | F value | Pr (>F) | Significance code |
|---|---|---|---|---|---|
| (Intercept) | 0.01 | 1 | 3.9996 | 0.045 | * |
| Subtype | 0.12 | 4 | 9.0704 | 0.000000254 | *** |
| Chromosome (Chr) | 0.46 | 22 | 6.247 | <2.2e-16 | *** |
| Ln(Distance) | 0.01 | 1 | 3.4926 | 0.062 | . |
| $\text{Ln(Distance)}^2$ | 0.01 | 1 | 3.8051 | 0.051 | . |
| $\text{Ln(Distance)}^3$ | 0.01 | 1 | 4.0679 | 0.044 | * |
| $\text{Ln(Distance)}^4$ | 0.01 | 1 | 4.291 | 0.038 | * |
| Subtype:Chr | 10.51 | 88 | 35.3659 | <2.2e-16 | *** |
| Subtype: Ln(D) | 0.13 | 4 | 9.689 | 7.83e-08 | *** |
| Chr: Ln(D) | 0.52 | 22 | 6.9751 | <2.2e-16 | *** |
| Subtype: $\text{Ln(D)}^2$ | 0.14 | 4 | 10.2293 | 0.000000028 | *** |
| Chr: $\text{Ln(D)}^2$ | 0.55 | 22 | 7.427 | <2.2e-16 | *** |
| Subtype: $\text{Ln(D)}^3$ | 0.15 | 4 | 10.7596 | 1.02e-08 | *** |
| Chr: $\text{Ln(D)}^3$ | 0.57 | 22 | 7.7224 | <2.2e-16 | *** |
| Subtype: $\text{Ln(D)}^4$ | 0.15 | 4 | 11.2843 | 3.73e-09 | *** |
| Chr: $\text{Ln(D)}^4$ | 0.59 | 22 | 7.9382 | <2.2e-16 | *** |
| Subtype:Chr: Ln(D) | 12.21 | 88 | 41.112 | <2.2e-16 | *** |
| Subtype:Chr: $\text{Ln(D)}^2$ | 13.74 | 88 | 46.2488 | <2.2e-16 | *** |
| Subtype:Chr: $\text{Ln(D)}^3$ | 15.07 | 88 | 50.7194 | <2.2e-16 | *** |
| Subtype:Chr: $\text{Ln(D)}^4$ | 16.22 | 88 | 54.5922 | <2.2e-16 | *** |
| Residuals | 981.23 | 290646 | | | |
